# Enzymatic and structural characterization of the *Naegleria fowleri* glucokinase

**DOI:** 10.1101/469072

**Authors:** Jillian E. Milanes, Jimmy Suryadi, Jan Abendroth, Wesley C. Van Voorhis, Kayleigh F. Barrett, David M. Dranow, Isabelle Q. Phan, Stephen L. Patrick, Soren D. Rozema, Muhammad M. Khalifa, Jennifer E. Golden, James C. Morris

## Abstract

Infection with the free-living amoeba *Naegleria fowleri* leads to life-threatening primary amoebic meningoencephalitis. Efficacious treatment options for these infections are limited and the mortality rate is very high (~98%). Parasite metabolism may provide suitable targets for therapeutic design. Like most other organisms, glucose metabolism is critical for parasite viability, being required for growth in culture. The genome of the parasite encodes a single glucose phosphorylating enzyme, a glucokinase (Glck). The products of this enzyme are required for both glycolysis and the pentose phosphate pathway. The *N. fowleri* Glck (NfGlck) shares limited (25%) amino acid identity with the mammalian host enzyme (HsGlck), suggesting that parasite-specific inhibitors with anti-amoeba activity could be generated. Following heterologous expression, NfGlck was found to have a limited hexose substrate range, with greatest activity observed with glucose. The enzyme had apparent *K*_m_ values of 42.5 ± 7.3 μM and 141.6 ± 9.9 μM for glucose and ATP, respectively. The NfGlck structure was determined and refined to 2.2 Å resolution, revealing that the enzyme shares greatest structural similarity with the *Trypanosoma cruzi* Glck. These similarities include binding modes and binding environments for substrates. To identify inhibitors of NfGlck, we screened a small collection of inhibitors of glucose phosphorylating enzymes and identified several small molecules with IC_50_ values < 1 μM that may prove useful as hit chemotypes for further lead and therapeutic development against *N. fowleri*.

## INTRODUCTION

Human infection by the free-living amoeba *Naegleria fowleri* can lead to life-threatening illness. When trophozoites encountered in freshwater are inadvertently introduced into the nasal passages, parasites can travel to the brain and cause a deadly infection, primary amoebic meningoencephalitis (PAM). Between 1962 and 2016, 143 PAM cases have been reported in the United States (Centers for Disease Control and Prevention). While the frequency of reported infection is low, the limited treatment options have historically yielded poor outcomes, with a study of 123 cases in the US revealing that 122 infections resulted in fatalities (1). A combination therapy that included amphotericin B, miconazole, fluconazole, and ketoconazole was used to successfully treat a single case of PAM, though the contribution of this therapeutic cocktail to patient survival is unclear (2). More recently, miltefosine has shown some promise as an anti-amoebic agent (3). Nevertheless, the lack of effective therapies for this infection, which remains lethal in ~99% of cases, makes elucidating therapeutic targets for novel drug discovery a high priority.

Mechanisms that the amoebae use to satisfy their metabolic needs are poorly resolved and are primarily based on assessment of growth under different culturing conditions and on analysis of the predicted gene content of the genome. The only *N. fowleri* metabolic enzyme characterized to date, phosphofructokinase (PFK), is a pyrophosphate-dependent (instead of ATP-dependent) enzyme (4). Use of the alternative phosphoryl group donor is usually associated with enzymes from anaerobic organisms, suggesting that *N. fowleri* may inhabit niches where oxidative phosphorylation is limited.

The role of glycolysis in meeting the metabolic needs of *N. fowleri* during human infection remains unresolved. However, the relative abundance of glucose in both the brain and cerebral-spinal fluid (CSF), and the correlation of parasite presence with reduced CSF glucose concentrations, suggest that glucose depletion in the brain may be a consequence of pathogen consumption (5). Therefore, the carbon source may be important for parasite metabolism.

Most eukaryotic cells rely on a hexose phosphotransferase (a hexokinase, HK) to catalyze the first enzymatic step common to both the glycolytic and the pentose phosphate pathways (PPP) to generate glucose-6-phophate (G6P). These enzymes typically have a broad affinity for different hexoses, including glucose, mannose, fructose, and galactose. *N. fowleri* harbors a single copy gene that encodes a predicted glucokinase (NfGlck) and lacks other recognizable enzymes that could catalyze the transfer of the γ-phosphoryl group of ATP to glucose to generate G6P. Glucokinases (Glck), which are typically restricted to using glucose as a substrate, have been classified into two groups, A and B (6). Group A Glcks include enzymes found in Gram-negative bacteria, Cyanobacteria, and amitochondriate protists. Group B Glck members include enzymes from Gram-positive bacteria that can use either ATP or polyphosphate as a phosphoryl group donor. Here, we describe the biochemical and structural characterization of NfGlck. This work has revealed that the enzyme has a limited hexose substrate use (like other Glcks) and shares structural similarity with the group A Glck from *Trypanosoma cruzi* (7). However, kinetic differences suggest the NfGlck may have evolved to satisfy the metabolic needs of *Naegleria* in low resource environments. Last, we describe the interrogation of a selected panel of potential inhibitors which has yielded the first NfGlck inhibitors disclosed to date. These compounds serve as a source of potential hits for further lead and therapeutic development.

## MATERIALS AND METHODS

### Chemicals and Reagents

Glucose-6-phosphate dehydrogenase, β-nicotinamide adenine dinucleotide phosphate (NADP^+^), and phosphoenol pyruvate (PEP) were purchased from Alfa Aesar (Haverhill, MA). β-nicotinamide adenine dinucleotide (NADH), adenosine triphosphate (ATP) and dimethyl sulfoxide (DMSO) were from Fisher Scientific (Pittsburgh, PA), while 2-phenyl-1,2-benzisoselenazol-3(2H)-one (ebselen, Eb, PubChem SID 856002) and glucosamine hydrochloride were obtained from VWR International (West Chester, PA). MMV688179 (2-[3-chloro-4-[5-[2-chloro-4-(diaminomethylideneamino)phenyl]furan-2-yl]phenyl]guanidine and MMV688271 (2-[2-chloro-4-[5-[3-chloro-4-(diaminomethylideneamino)phenyl]furan-2-yl]phenyl]guanidine) were provided by Medicines for Malaria Venture (Switzerland). The benzamidobenzoic acid derived compounds (entries 5-23, Table 1) were provided by Dr. Jennifer E. Golden at the University of Wisconsin-Madison with > 95% purity as determined by UV-MS. See supporting information for synthetic details and characterization.

**Table 1.** Crystallography parameters

|  |  |
| --- | --- |
| **Crystal parameters** |  |
| Space group | *I*422 |
| Cell dimensions<br>a=b=c (Å), $\alpha=\beta=\gamma$ | 202.48, 202.48, 68.79,<br>90, 90, 90 |
| **Data set** |  |
| X-ray Source | Rigaku F-RE+SuperBright |
| Wavelength (Å) | 1.5418 |
| Resolution (Å) | 50-2.20 (2.26-2.20) |
| Rmerge | 0.090 (0.572) |
| I/sigma (I) | 17.95 (3.27) |
| CC (1/2) | 99.0 (86.8) |
| Completeness | 99.6% (99.7%) |
| # reflections overall | 258,457 (13,992) |
| # reflections, unique | 36,362 (2,642) |
| Multiplicity | 7.1 (5.3) |
| **Refinement statistics** |  |
| Rwork | 0.1539 |
| Rfree | 0.1872 |
| RMSD bond lengths (Å) | 0.007 |
| RMSD bond angles (°) | 0.822 |
| Ramachandran: | - |
| favored (%) | 97.61 |
| allowed (%) | 2.39 |
| Outliers | 0 |
| Molprobity clash score | 1.46 |
| Molprobity score | 1.03 |
| PDB code | 6da0 |

### Growth Inhibition Assay

*N. fowleri* strain 69 (ATCC 30215) trophozoites were cultured axenically at 37°C in Nelson’s Complete Media (NCM, 0.17% liver infusion broth (BD Difco, Franklin Lakes, NJ), 0.17% glucose, 0.012% sodium chloride, 0.0136% potassium phosphate monobasic, 0.0142% sodium phosphate dibasic, 0.0004% calcium chloride, 0.0002% magnesium sulfate, 10% heat inactivated fetal bovine serum (FBS), 1% penicillin-streptomycin) in tissue culture treated flasks (Corning 353135). Trophozoties were seeded at 2 x 10^6^ cells/mL in 10 mL of NCM, or NCM lacking glucose and cultures were imaged daily on an EVOS Imaging System (Thermo Fisher Scientific, Waltham, MA).

### Recombinant NfGlck Purification

The open reading frame NF0035880 for the *N. folweri* glucokinase (NfGlck) (Amoebadb.org) was synthesized for codon optimization (Genescript, Piscataway, NJ), sequenced, and cloned into pQE-30 (Qiagen, Valencia, CA).

Recombinant NfGlck, a ~49 kDa protein, was purified following a protocol developed for heterologous expression and purification of a HK from the African trypanosome. Briefly, a 8 mL bacterial culture of *E. coli* M15(pREP4) harboring pQE30-NfGlck was used to inoculate a 4 L culture which was grown to an OD_600_ of ~0.6 and then induced for 24 h at room temperature with 500 μM isopropyl β-D-1-thiogalactopyranoside (IPTG) and purified using an approach modified from (8). After nickel column isolation (HisTrap FF, GE Healthcare Life Sciences, Pittsburgh, PA), active fractions were dialyzed into 20 mM Tris-HCl pH 8, 50 mM NaCl and further purified by anion exchange chromatography on a HiTrap Q XL (GE Healthcare Life Sciences, Pittsburgh, PA). The purified protein was stored in 25 mM HEPES-Na pH 7.4, 150 mM NaCl and 50% glycerol. Protein concentration was determined by Bradford assay using BSA to generate a standard curve.

### Glucokinase Assay

Enzyme was assayed in triplicate using a coupled reaction to measure enzyme activity. Briefly, enzyme (10 nM) in assay buffer (50 mM TEA-HCl, 3.3 mM MgCl_2_, pH 7.4) was added clear 96-well plates and incubated for 15 minutes at room temperature. To start the reaction, 100μL of substrate buffer (425 μM glucose, 1.5 mM ATP, 1mU/μL G6PDH, and 0.75 mM NADP^+^) was added to each well. The absorbance at 340 nm was then measured on a Biotek Synergy H1 microplate reader every 15 seconds for 2 minutes. Alternatively, enzyme activity was measured in a coupled reaction containing pyruvate kinase and lactate dehydrogenase. Briefly, the 200 μL reaction was performed in 96-well plates with NfGlck (10 nM) incubated in reaction buffer containing 50 mM TEA-HCl pH 7.4, 3.3 mM MgCl_2_, 20 mM Hexose, 2 mM ATP, 2.5 mM PEP, 0.6 mM NADH and 1 μL pyruvate kinase-lactate dehydrogenase (Sigma, St. Louis, MO). This reaction measures the production of ADP through the coincident change in absorbance at 340 nm. Kinetic analyses were performed using KaleidaGraph 4.1 (Synergy Software, Reading, PA).

### Expression and purification of NfGlck for crystallography

The gene for NfGlck (AmoebaDB NF0035880) was amplified from *N. fowleri* RNA (the generous gift of Dr. Dennis Kyle, the University of Georgia) and cloned into the expression vector pBG1861 using ligand-independent cloning (9). The expression vector provides a non-cleavable N-terminal His6-tag (SSGCID target ID NafoA.19900.a, SSGCID construct ID NafoA.19900.a.B11, SSGCID batch NafoA.19900.a.B11.PW38443). NfGlck was expressed in *E. coli* Rosetta BL21(DE3)R3 following standard SSGCID protocols as described previously (10). Purification was done using Ni-NTA affinity and size exclusion chromatography following standard SSGCID protocols (11). The purified protein was concentrated to 13.78 mg/mL in its final buffer (25mM HEPES pH 7.0, 500mM NaCl, 5% glycerol, 2mM DTT, 0.025% NaN3), flash frozen in liquid nitrogen and stored at −80°C.

### Crystallization, Data collection and Structure solution

Crystallization set ups were performed using the apo protein and with the addition of 2 mM each of MgCl_2_, AMPPNP, and D-glucose, and NfGlck at 13.78 mg/mL in 96-well XJR crystallization trays (Rigaku Reagents) with 0.4 μL protein mixed with 0.4 μL reservoir, equilibrating against 80 μL reservoir solution. Crystallization conditions were searched for by using sparse matrix screens JCSG+ (Rigaku Reagents), MCSG1 (Microlytic/Anatrace) and Morpheus (Molecular Dimensions). Crystallization trays were incubated at 285 K. Several crystals were observed in the ligand trays, while no crystals grew in the apo trays.

Crystals from Morpheus condition C8 (12.5% w/v PEG 1000, 12.5% w/v PEG 3350, 12.5% v/v MPD; 30 mM of each sodium nitrate, disodium hydrogen phosphate, ammonium sulfate; 100 mM MOPS/HEPES-Na pH 7.5) were harvested and vitrified without adding additional cryo-protectant by plunging the crystals in liquid nitrogen. A diffraction data set was collected in-house on a Rigaku FR-E^+^ SuperBright rotating anode equipped with Rigaku VariMax optics and a Saturn 944+ detector, using CuKα X-rays. The diffraction data were reduced with the XDS package (12), and are summarized in Table 1.

The structure was solved using the program MorDa (13) from the CCP4 package (14). The initial model was refined and extended using the phenix.ligand_pipeline script within Phenix (15). Manual model building was done with Coot (16) and the structure was refined in reciprocal space with Phenix.refine (15). Ligand restraints for AMPPNP and β-*D*-glucose were generated using the Grade web server (http://qrade.qlobalphasinq.org). The structure was validated using the built-in validation tools within Phenix. The coordinates and structure factors were deposited in the PDB with accession codes 6DA0.

## RESULTS

The dietary requirements of the free-living amoeba *Naegleria fowleri* are poorly defined, with parasite growth described from cultures maintained in the presence of bacteria or human feeder cells. Axenic culture in defined media has also been established (17), but the role of individual components in the metabolic success of the parasites has not been fully explored.

Most of the described media includes glucose. To assess the importance of this carbon source to the parasite, trophozoites were seeded into standard NCM with or without glucose (Fig.1A). After two days, culture in the absence of glucose led to reduced growth and enhanced encystation. These cultures began to encyst earlier than cells cultured in the presence of glucose and as a consequence failed to reach the density of cells under standard conditions. To further consider the importance of glucose to the parasites, trophozoites were incubated with the glycolytic poison 3-bromopyruvate (3BP) in standard glucose-bearing (9 mM glucose) medium. Inclusion of the compound, which is known to inhibit the glycolytic enzyme glyceraldehyde-3-phospate dehydrogenase (GAPDH) (18) led to reduced cell growth (Fig. 1B), with only cysts detectable after two days of incubation.

**Fig. 1.**
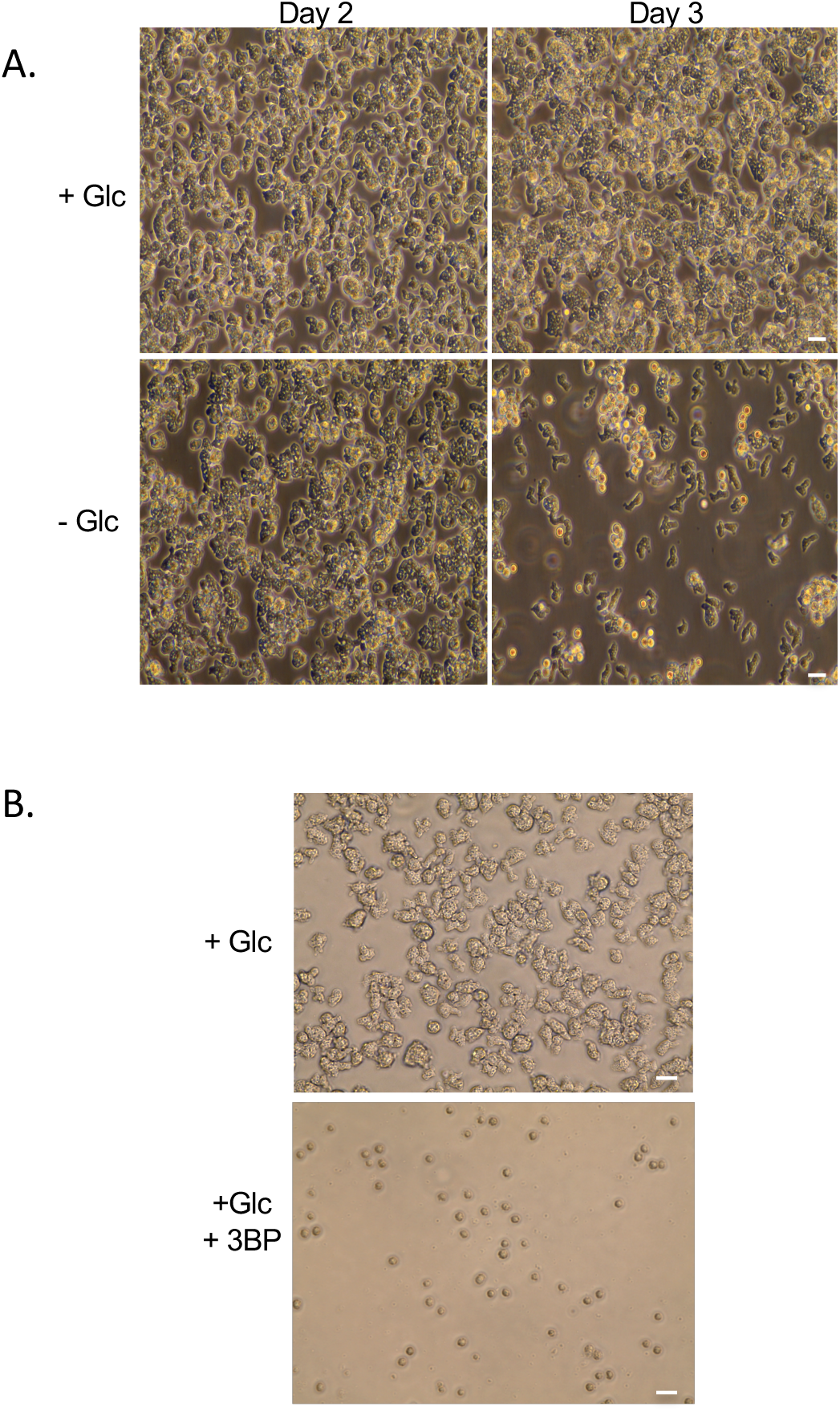
(A) *N. fowleri* trophozoites were seeded at 2 x 10^6^ cells/mL in control media (Nelson’s Complete Media) and cultured with or without glucose. (B) Cells (1 x 10^6^ cells/mL) were incubated with or without 3-bromopyruvate (50 mM) for two days prior to imaging. Images were captured on an EVOS Imaging System and data is representative of multiple experiments. Scale bar = 20 μM.

The absence of glucose or treatment with 3BP could be impacting *N. fowleri* by interfering with metabolism (through glycolysis) or sugar nucleotide biosynthesis (through the pentose phosphate pathway). Entry into either of these pathways requires glucose-6-phosphate, which is typically generated by HK or Glck enzymes, suggesting inhibitors of this activity could be useful anti-amoebic compounds. The *N. fowleri* genome does not encode obvious HK-like enzymes but does harbor a single gene for a putative Glck. Unlike HKs, which can transfer a phosphoryl group to different hexoses, Glcks have a restricted substrate range, primarily utilizing glucose as an acceptor for the γ-phosphoryl of ATP.

Beyond Glcks from other *Naegleria spp*. and related amoebae, NfGlck shares limited identity (37%) with the Glck from *Trichomonas vaginalis*, a protist that lives primarily under anaerobic conditions. The enzyme was less similar to HsGlck, *T. cruzi* Glck (TcGlck), and *Leishmania donovani* Glck, with sequence identities of 25%, 27%, and 26%, respectively.

Recombinant NfGlck, which was purified to ~99% homogeneity (as determined by Coomassie staining of an SDS-PAGE gel), was determined to be a monomer by gel filtration chromatography. The enzyme displayed Michaelis-Menten kinetics when glucose levels were increased (Fig. 2A) and had an apparent *K*_m_ value of 42.5 ± 7.3 μM for the sugar, a *k*_cat_ value of 36.5 sec^-1^, and a pH optimum of 8.5 (Fig. 2C). NfGlck had a limited number of sugar substrates it could use in catalysis, with detectable activity when glucose, glucosamine, or fructose (at high concentration) were used as sugar substrates (Fig. 3A). Glucosamine was also a competitive inhibitor of glucose, with a *K*_i_ of 186.4 ± 20.6 μM. The enzyme had an apparent *K*_m_ value of 141.6 ± 9.9 μM for ATP (Fig. 2B). While AMP and PP_i_ had no impact, ADP was an inhibitor of the enzyme, with a *K*_i_ of 116.3 ± 7.0 μM (Fig. 3B). The enzyme functioned best when MgCl_2_ was included in the assay with MnCl_2_ and CaCl_2_ yielding 40% and 9%, respectively, of the activity observed when MgCl_2_ was included in the assay.

**Fig. 2.**
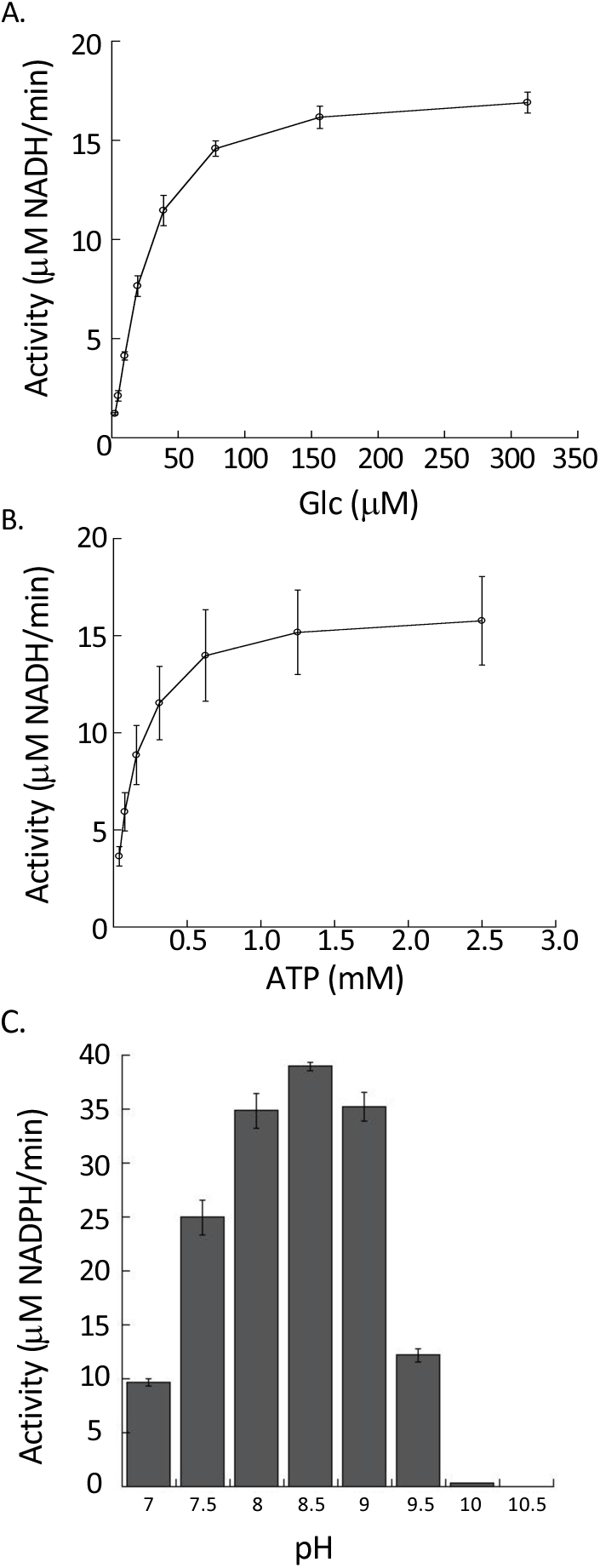
Effects of ATP, glucose, and pH on NfGlck activity. (A) NfGlck (10 nM) activity in response to increasing (A) glucose (B) ATP concentrations or to different (C) pH levels. Two different buffers with buffering capacity in the range of pH tested were used for the assay: Tris-Cl (pH 7-8.5) and sodium carbonate (pH 9.0-10.5). Reactions were carried out in triplicate, and standard deviations are indicated.

**Fig. 3.**
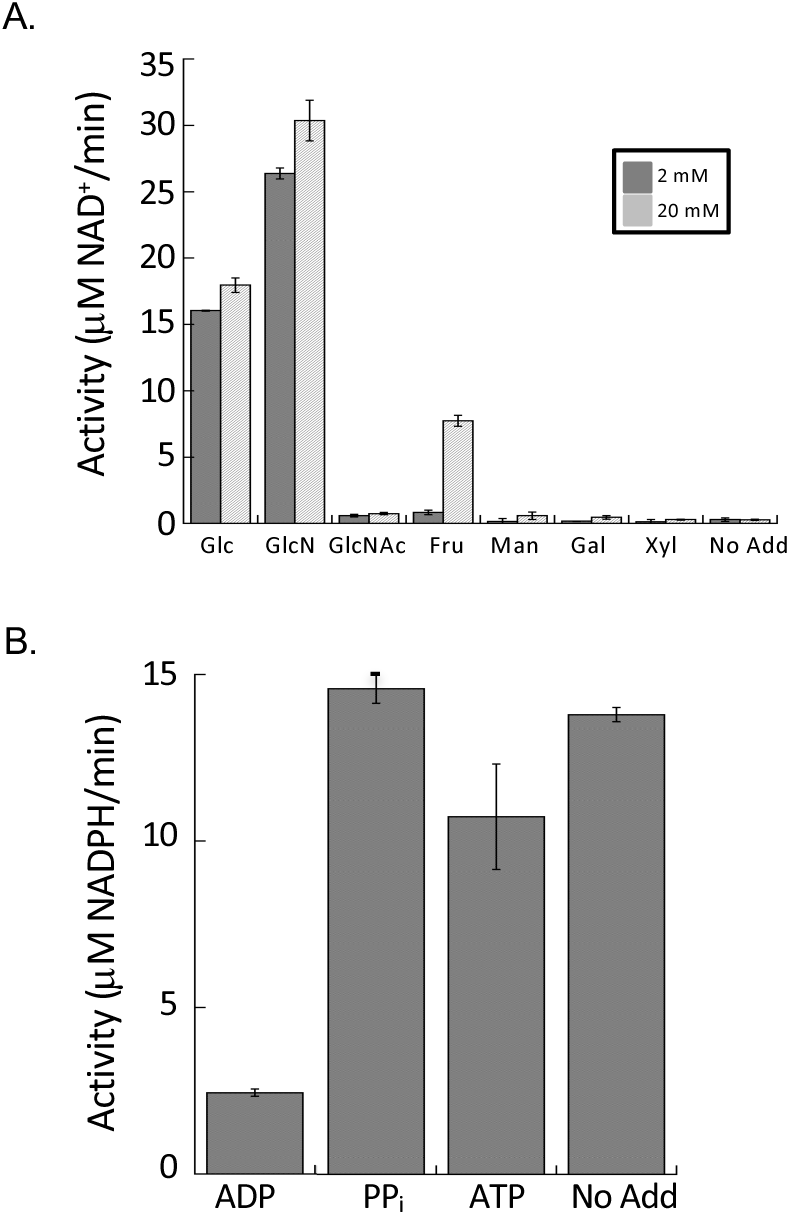
(A) NfGlck (10 nM) activity with various hexoses (5 mM). (B) Impact of phosphoryl bearing substrates on NfGlck activity. NfGlck activity was measured in presence of 1.5 mM ATP supplemented with ADP, PP_i_, and ATP (all at 5 mM). Assays were carried out in triplicate and standard deviations are indicated.

### Structural analysis of NfGlck

The structure of NfGlck was solved by Molecular Replacement despite low sequence homology with structures in the PDB. The closest homolog by sequence in the PDB is the *T. cruzi* Glck (TcGlck) with 27% sequence identity. This protein is also the source of the closest structural homologs in the PDB, which are structures of the TcGlck in complex with β-*D*-glucose and ADP (PDB code 2Q2R, 1.7Å RMSD), in complex with CBZ-GLCN (5BRE, 1.8Å RMSD), in complex with BENZ-GLCN (5BRD, 2.0Å RMSD), HPOP-GLCN (5BRF, 2.0Å RMSD), and DBT-GLCN (5BRH, 2.0Å RMSD) (19). The structure of HsGlck (3FGU) superimposes with an RMSD of 3.0Å. In each of the alignments, ~300 out of the 370-380 residues were aligned. Structure solution was facilitated by the domain approach of MorDa (13), which placed three domains from PDB entries 5BRH, 1SZ2 and 3AAB with an overall RMSD of 1.2Å for 282 aligned Cα atoms. HsGlck shares little sequence homology with NfGlck. The closest structural homolog is HsGlck in complex with an allosteric inhibitor (4MLH), with an RMSD of 2.9Å over 244 residues. While NfGlck and HsGlck share a similar fold and glucose binding domain, the noted sequence and structural diversity between the two enzymes suggests that specific inhibitors of NfGlck could be developed.

The structure of NfGlck was refined against diffraction data up to 2.2Å resolution (Fig. 4). The structure refines with appropriate quality indicators (Table 1) and NfGlck crystallizes with one molecule per asymmetric unit. N-terminal residues Met1-Glu23, and C-terminal residues Gly401-Lys443 do not have the sufficient electron density needed to be modeled, while a consecutive model could be built for residues Pro23-Leu400. NfGlck consists of two domains, a small α/β domain (residues N-Asn171 and Asn384-C) and a larger α/β domain (His174-Glu382). NfGlck was crystallized in the presence of AMPPNP and D-glucose and both molecules could be modeled with confidence, even though AMPPNP did not appear to be fully occupied (81% refined occupancy). While D-glucose is located at the domain interface, AMPPNP mostly binds to the large domain.

**Fig. 4.**
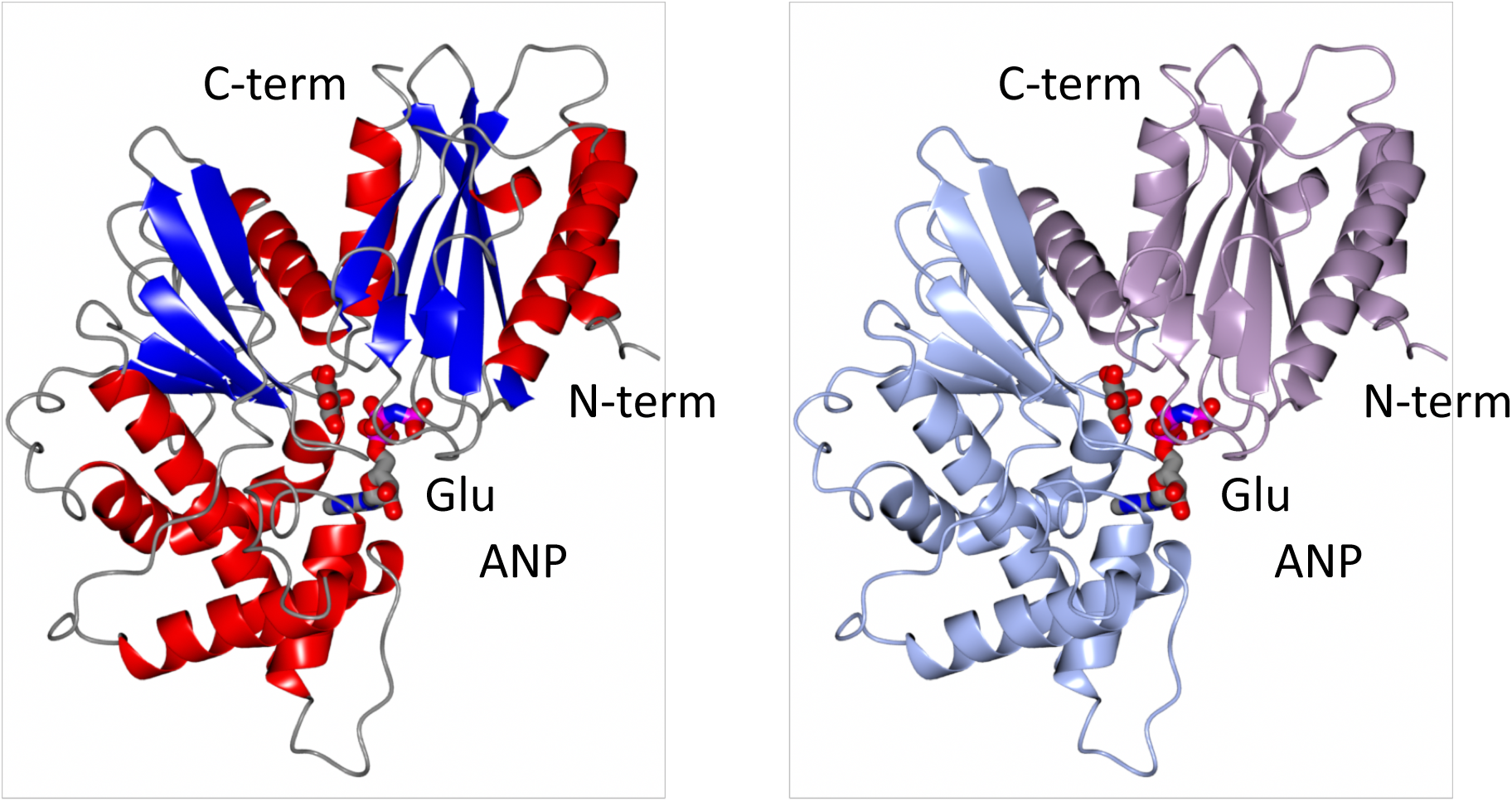
The structure of NfGlck bound to AMPPNP and glucose is shown in two color schemes. The left structure is colored by secondary structure elements. The right structure shows the smaller α/β domain (N-Asn171 and Asn384-C) in lavender, and the larger α/β domain (His174-Glu382) in light blue. N- and C-termini are indicated and glucose and AMPPNP are the left and right ball-and-stick represented ligands.

Despite the low sequence identity and high RMSDs, NfGlck and TcGlck have an identical topology; all secondary structure elements match up well. Notable exceptions are loops around residues Leu42-Phe47, Pro179-Gln190 and Gly238-247. None of these loops are involved ligand binding. NfGlck appears to adopt the same nearly-closed conformation as was described for inhibitor-bound TcGlck structures. The binding modes and binding environment for both D-glucose and AMPPNP or ADP, respectively, in NfGlck and TcGlck (2QRQ) are also very well conserved. In TcGlck, there is no equivalent to the hydrogen bond of Thr127 with the 1-hydroxyl group of glucose. Similarly, in NfGlck, His231 provides additional hydrogen bonds with the 1- and 2-hydroxyl groups of glucose. This interaction is not available in TcGlck where the equivalent Ser210 does not reach the glucose (Fig. 5A and B). The amine moiety of glucosamine would extend towards His231. This might explain the different specificities of NfGlck and TcGlck for gluosamine as a substrate. In HsGlck, the glucose binding is a blend of the NfGlck and the TcGlck binding mode: There is no direct replacement for the interaction of His231 (Fig. 5D). However, in HsGlck Thr168 has a similar role as Thr127 in NfGlck.

**Fig. 5.**
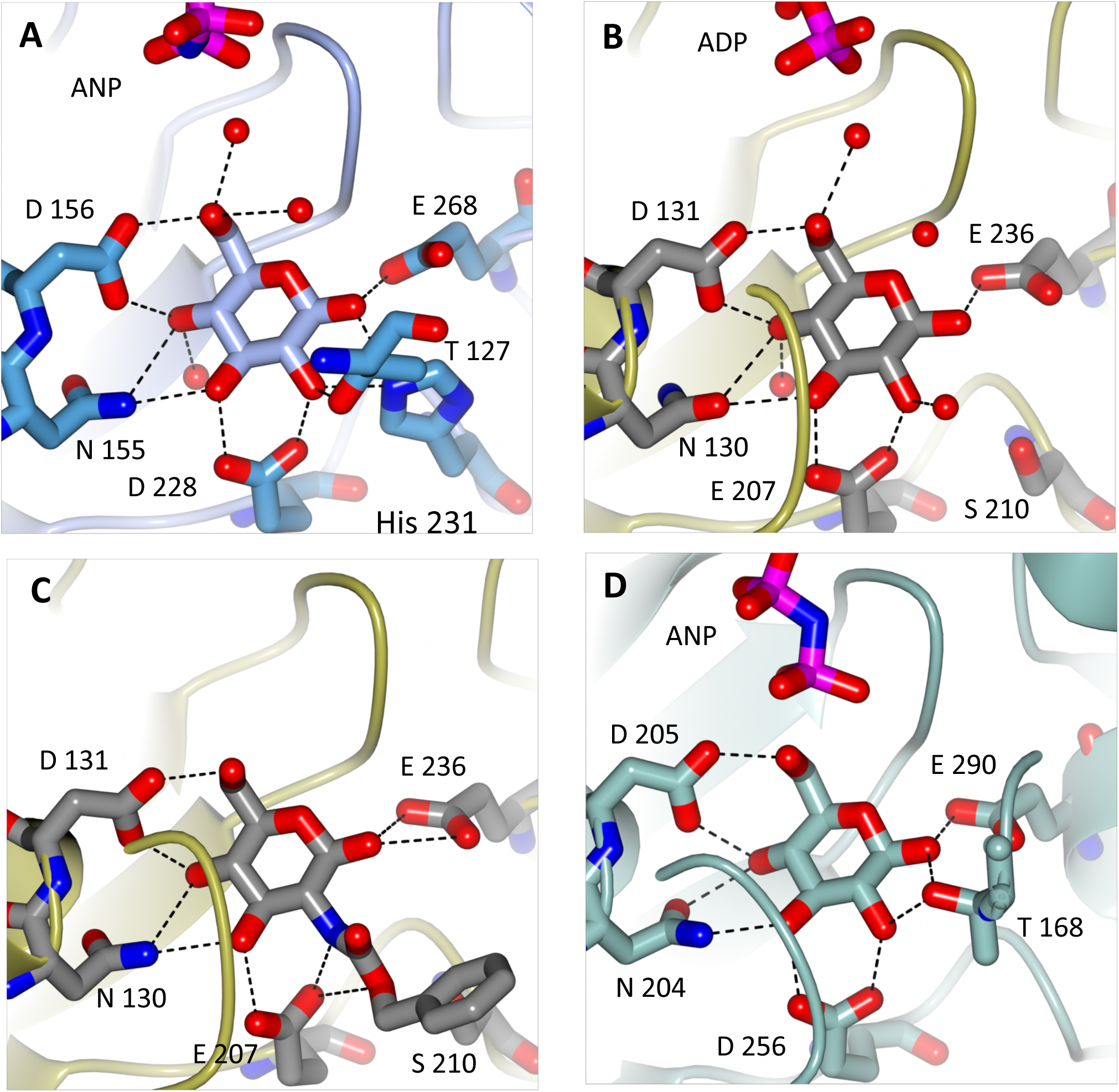
Glucose binding sites of various Glcks. (A) In NfGlck, glucose is bound with a strong hydrogen bond network, including hydrogen bonds from the 1- and 2-hydroxyl groups to His231. In TcGlck (B) glucose is bound with a very similar hydrogen bond pattern. Ser210 replaces His231 and cannot engage in a hydrogen bond with glucose. The space that His231 occupies in NfGlck is the part of the glucosamine moiety of glucosaminelike inhibitors of TcGlck (C). In HsGlck (D), most of the hydrogen bond network is conserved, with the 1- and 2-hydroxy groups hydrogen bound to Thr168 from a different part of the protein.

Additional TcGlck structures have been described in the presence of several glucosamine analogue inhibitors (Fig. 5C). While the glucose moiety of the ligand binds identically to glucose, the region that the substituent of the ligands occupies in TcGlck is significantly different in NfGlck. Most notably, Phe339 in TcGlck is Trp370 in NfGlck which would provide a steric hindrance for the ligands. Phe337, which provides a significant hydrophobic interaction in TcGlck is His369 in NfGlck. It would require a conformational change in NfGlck compared with the presented crystal structure for His369 to engage in this hydrophobic interaction. In TcGlck, neither Phe337 nor Phe339 shows very distinct conformational differences between the ADP/Glucose and the glucosamine analogues structures. By extension, these two residues are expected to provide a very different binding profile of glucosamine analogue inhibitors for NfGlck and TcGlck.

### Identification of small molecule inhibitors of NfGlck

To identify inhibitors of NfGlck, a subset of known HK inhibitors and related structural analogs were tested against the *N. fowleri* glucokinase enzyme. For example, the screening collection assessed for this effort contained compounds selected from the Pathogen Box chemical library that were known to inhibit *T. brucei* hexokinase 1 (TbHK1), other structurally distinct small molecule inhibitors identified in our TbHK1-based HTS campaigns, and analogs subsequently developed from these hits (20). The advantage of using these compounds included the likelihood of inhibition of a related parasitic metabolic enzyme and, due to our past experiences with validating these compounds, pre-existing data that showed (a) that these agents did not interfere with the coupled enzyme used in the screen and (b) that the compounds were generally non-toxic to mammalian cells (MMV, (20)).

NfGlck was inhibited by two structurally-related bis-guanadinyl compounds from the Pathogen Box collection, MMV688179 and MMV68827, which had anti-parasitic activity against African trypanosomes (21). Interestingly, MMV688179 yielded an IC_50_ value of 2.02 ± 0.31 μM against NfGlck and was similarly active against TbHK1 and PfHK with IC_50_ values of 2.90 ± 1.13, and 0.78 ± 0.17 μM, respectively. This compound, which was a mixed inhibitor with respect to ATP, did not inhibit HsGlck at 10 μM (Table 2). Comparatively, the related compound MMV688271 was a mixed inhibitor with respect to ATP and showed modestly more potent inhibition of parasite enzymes with IC_50_ values at least 2-fold better than MMV688179. However, the two compounds differed in that MMV688271 was less selective, inhibiting HsGlck with an IC_50_ value of 2.58 ± 0.53 μM, a value similar to those found for the parasite enzyme inhibition. Neither Pathogen Box compounds were toxic to *Naegleria* at 10 μM, but given their IC_50_ values, this was not surprising (data not shown).

**Table 2.**
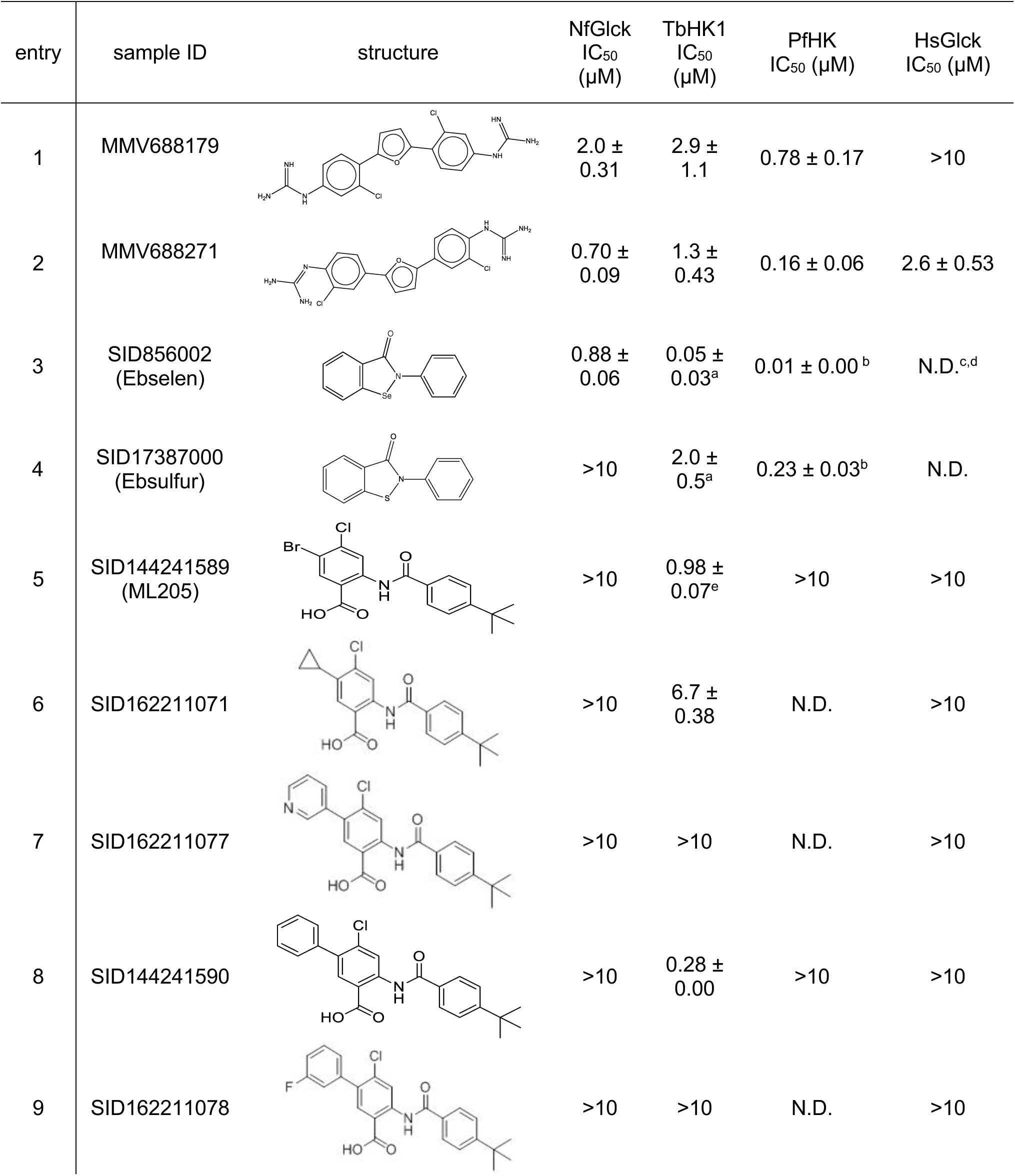
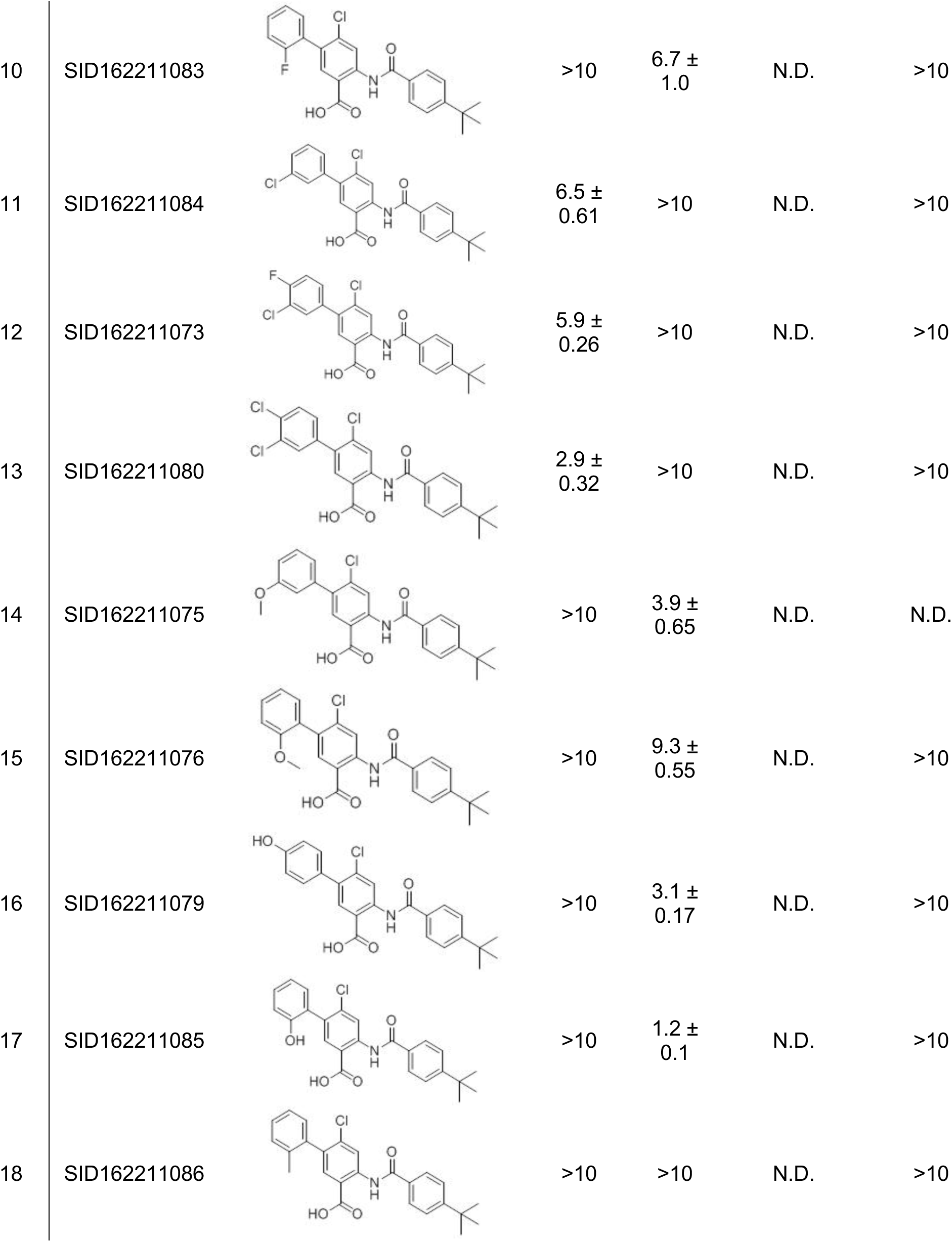
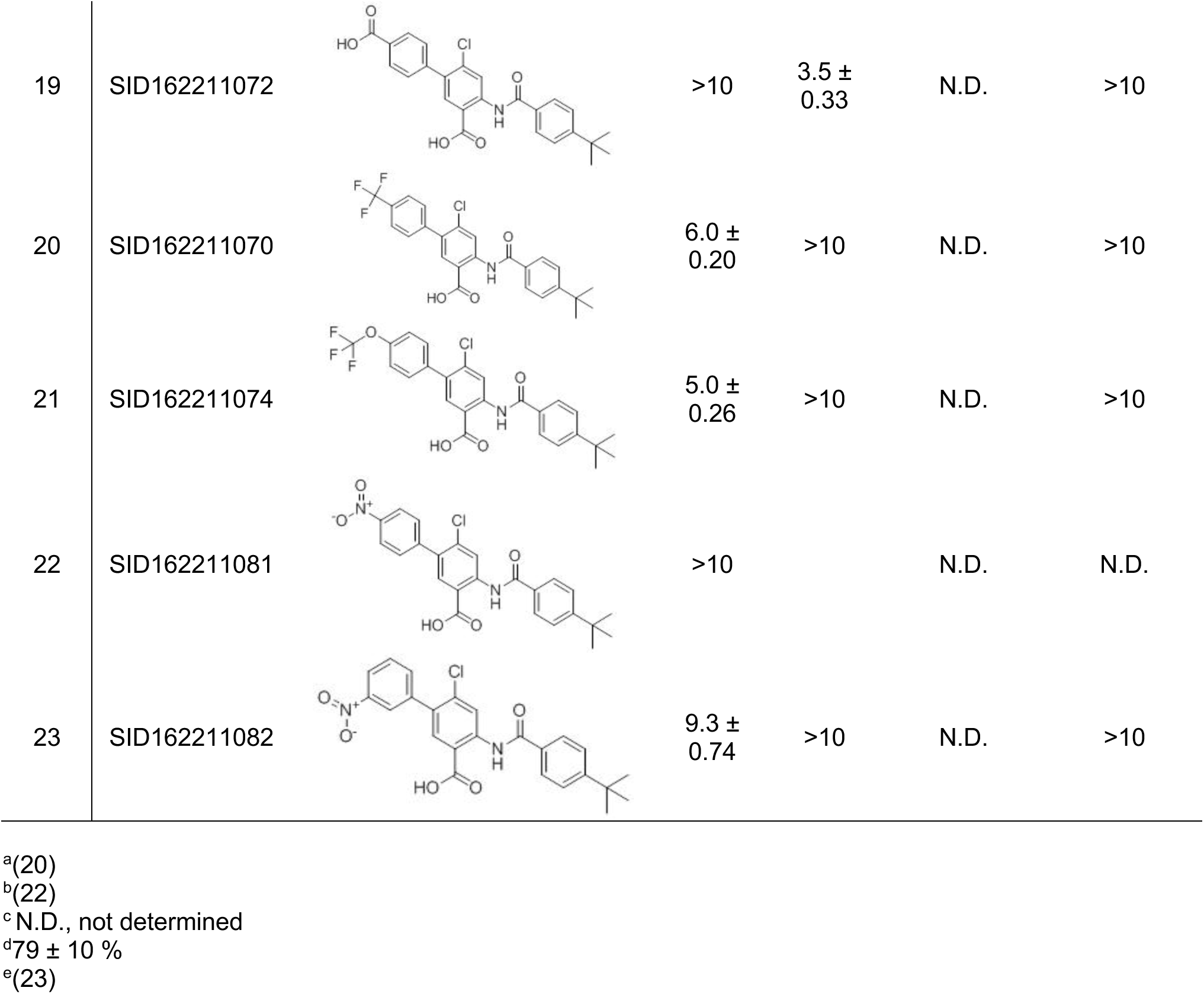
Small molecule inhibitors of NfGlck

From the TbHK1 HTS, two distinct scaffolds were identified and pursued for analog development against TbHK1. Ebselen (SID856002) and the structurally similar isobenzothiazolinone (SID 17387000) were identified as potent inhibitors of the trypanosome enzyme and are active against the *Plasmodium* HK (22). Of the two agents, only ebselen showed inhibition of NfGlck with an IC_50_ of 0.88 μM, a notably higher concentration than that required to inhibit the other parasite HKs (Table 2). The second scaffold, a benzamidobenzoic acid-based inhibitor of TbHK1 (20), was developed through structure-activity relationship (SAR) optimization against the trypanosome to yield ML205, a potent inhibitor of TbHK1 (23). While this particular compound was not active against the *Naegleria* enzyme (IC_50_ > 10 μM), evaluation of related analogs revealed some promising activity.

Analogs of ML205 tested in this work included compounds featuring substitution of the bromine atom on the benzoic acid core. One of the best modifications found for inhibition of TbHK1 involved exchange of the bromine atom of ML205 with a phenyl ring to afford SID144241590 with a TbHK1 IC_50_ = 0.28 μM; however, this analog showed no significant inhibition of NfGlck. Likewise, bromine substitution with a cyclopropyl group (SID162211071), a 3-pyridyl moiety (SID162211077), or mono-fluorinated phenyl groups (SID162211078 and SID162211083) did not generate analogs that inhibited NfGlck, despite showing submicromolar activity in some cases against TbHK1.

While mono-fluorination of the phenyl appendage did not impact NfGlck, incorporation of chlorinated phenyl rings was advantageous against this enzyme. The analog bearing a 3-chlorophenyl group (SID162211084) gave moderate NfGlck inhibition (IC_50_ = 6.5 μM), the 3-chloro-4-fluorophenyl analog (SID162211073) trended with marginally better potency, and the 3,4-dichlorophenyl derivative (SID162211080) showed the most improved potency against NfGlck with an IC_50_ = 2.9 μM. Alternative phenyl ring substitutions, which included hydroxylation, methyl or methoxy groups, did not afford analogs with appreciable inhibition of NfGlck, although several of these did demonstrate reasonable inhibition of TbHK1. Other analogs that inhibited NfGlck with potency better than 10 μM featured a 4-trifluoromethylphenyl or 4-trifluoromethoxyphenyl group (SID162211070 and SID162211074, respectively) or a 3-nitrophenyl group (SID162211082). Relative to ML205, the data suggest that elaborated phenyl rings in place of the bromine atom help to refine potency against NfGlck, and the specific substitution patterns that are favored are a result of a combination of spatial and electron-withdrawing effects. Further, this effort revealed that the structural elements that improve NfGlck inhibition appear to differ from those driving TbHK1 inhibition for this particular structural series as it relates to changes in this region of the scaffold. For instance, analogs with electron-donating substituents on the phenyl ring registered with some TbHK1 inhibition but did not do the same for NfGlck. Alternatively, analogs with more electron-deficient phenyl rings appended to the core favored NfGlck inhibition and not TbHK1.

## DISCUSSION

As a thermophilic free-living amoeba, *N. fowleri* is found world-wide in or nearby warm freshwater. In the freshwater habitat, free glucose in the water column is likely limited thus raising the possibility that NfGlck has only a limited role in parasite metabolism in this environment. However, the amoeboid trophozoite feeds on a variety of prey in the water including bacteria, fungi, and other protists, which may generate substrates for NfGlck. The sediment at the bottom of freshwater sources is different, with free sugars including glucose (1.1 mg/g dry weight) characterized in lake sediment samples (24, 25). Carbohydrates have also been characterized from lake surface mud samples, with glucose found at 13-191 mg/kg dry weight (26). These findings suggest that NfGlck may play a role in metabolism of nutrients acquired from the environment through either predation or directly from the sediment.

What role could NfGlck play during human infection? The process is usually initiated following forceful introduction of water contaminated with trophozoites into the nasal cavity. The presence of the trophozoites induces the host to secrete mucus, a response that serves as a barrier to invasion of the nearby tissue (27). The mucus contains an abundance of glycoproteins (including mucins), which compose the majority of its dry weight (28). To overcome this obstacle, the parasites release enzymes including glycosidases that degrade the mucin glycan components, potentially liberating substrates for NfGlck while aiding the parasite in accessing underlying tissue for ultimate invasion of the brain (29).

Degradation of host tissues during invasion provides access to glucose and glucose-bearing host macromolecules. As the parasite attacks the host cribriform plate, glucose availability is presumably enhanced once sufficient tissue damage has occurred to allow access to host blood, which has glucose levels that are maintained at ~5 mM. The brain infection that follows exposes the parasite to lower (though still abundant) glucose concentrations and the presence of parasites has been implicated in reducing the overall CSF glucose levels (5).

Whether NfGlck plays a central role in glucose catabolism in the diverse environments in which the parasite can reside remains to be determined. However, because the enzyme also provides substrate to the pentose phosphate pathway (PPP) for the generation of sugar nucleotides, the activity is likely indispensable, a possibility that awaits the development of suitable molecular tools for genetic ablation of the enzyme activity *in vivo*. Nevertheless, the position of the enzyme at the head of glycolysis and the PPP suggests that interfering with its activity could yield potent anti-parasitic compounds, and the differences in sequence and structure of NfGlck and HsGlck suggest inhibitor specificity could be established. Here, we have identified a number of enzyme inhibitors but none impaired parasite growth at the levels tested (typically 10 μM). This may be for various different reasons, including abundant substrate competitor molecules in the cytoplasm and insufficient access to the target in the cell. It is likely that much more potent inhibitors are required to see inhibition of parasite growth.

The NfGlck structure is highly similar to that of TcGlck. The *T. cruzi* enzyme was noted for having a structure consistent with that described for *E. coli* glucokinase (EcGlck), a group A Glck, which is a member of the group A ribokinase enzymes (19). Like other group A enzymes, NfGlck was limited to using ATP as a phosphoryl donor. However, kinetic differences between NfGlck and TcGlck raise the possibility that the *Naegleria* enzyme is an atypical group A member. This is primarily based on the observation that NfGlck, which can use glucose, glucosamine, or fructose as a substrate, lacks the glucose-limited substrate specificity that is a hallmark of group A enzymes. NfGlck and TcGlck also have distinct kinetic properties (7). NfGlck has an affinity for glucose that is ~23-fold higher than TcGlck, perhaps explained by the additional hydrogen bonds provided by NfGlck His231 with the 1- and 2-hydroxyl groups of glucose, which are not present in TcGlck where the equivalent residue is a Ser (Fig. 5A and B). Additionally, ADP inhibited NfGlck at much lower concentrations than those required for TcGlck inhibition (7).

The finding that NfGlck requires ATP for activity seems at odds with the presence of a *N. fowleri* PFK that is PP_i_-dependent (4). Other pathogenic protists, like the anaerobes *Entamoeba histolytica* and *Trichomonas vaginalis*, have PP_i_-dependent glycolytic enzymes (4, 30), presumably because the alternative phosphoryl group donor allows optimized ATP production under energy-limiting conditions. The metabolic flexibility noted in *Naegleria gruberi* supports the notion that *N. fowleri* may have a PP_i_-dependent PFK in order to support anaerobic fermentation (31). *N. fowleri* would likely encounter oxygen-poor environments, particularly when lake sediments are at higher temperatures during summer months. Alternatively, it is possible that an ATP-dependent PFK exists and has not yet been detected. It is known, for example, that some organisms can express either an ATP-dependent or PP_i_-dependent PFK in response to the types of environmental carbon sources available (32). Lastly, in some organisms a PP_i_-dependent PFK has been postulated to function as both a PFK in glycolysis and a fructose-bisphosphatase in gluconeogenesis (33). This is possible because thermodynamically (and under physiological conditions) PP_i_ can be either consumed or regenerated. A similar role for the *N. fowleri* PP_i_-dependent PFK is unlikely, as parasites do not grow in the absence of glucose (Fig. 1). This suggests that any G6P generated by the PP_i_-dependent PFK is insufficient to meet the metabolic demands of the parasite, and supports the notion that NfGlck is essential.

## ACKNOWLEDGMENTS

The authors would like to thank Dr. Dennis Kyle for providing the *N. fowleri* RNA library (to SSGCID). Work from the JCM laboratory was supported in part by the US National Institutes of Health Center for Biomedical Excellence (COBRE) grant under award number P20GM109094. This project has also been funded in part with Federal funds from the National Institute of Allergy and Infectious Diseases, National Institutes of Health, Department of Health and Human Services, under Contract No.: HHSN272201700059C. JEG acknowledges institutional support from the Office of the Vice Chancellor for Research and Graduate Education at the University of Wisconsin-Madison with funding from the Wisconsin Alumni Research Foundation (WARF): UW2020 225 infrastructure grant (JEG, PI). This work made use of the instrumentation at the UW-Madison Medicinal Chemistry Center, funded by the UW School of Pharmacy. MK was supported by the John & Jane Roudebush Distinguished Graduate Fellowship (School of Pharmacy Pharmaceutical Sciences Division) and the Daniel H. Rich Distinguished Scholarship and the National Science Foundation Graduate Research Fellowship Program under Grant No. DGE-1256259.

## The abbreviations used are

G6P: glucose-6-phosphate
G6PDH: glucose-6-phosphate dehydrogenase
HK: hexokinase
HsGlck: *Homo sapiens* glucokinase
NfGlcK: *Naegleria fowleri* glucokinase
PfHK: *Plasmodium falciparum* hexokinase
TbHK1: *Trypanosoma brucei* hexokinase 1
TcGlck: *Trypanosoma cruzi* glucokinase

